# FlatProt: 2D visualization eases protein structure comparison

**DOI:** 10.1101/2025.04.22.650077

**Authors:** Tobias Olenyi, Constantin Carl, Tobias Senoner, Ivan Koludarov, Burkhard Rost

## Abstract

**Background:** Understanding and comparing three-dimensional (3D) structures of proteins can advance bioinformatics, molecular biology, and drug discovery. While 3D models offer detailed insights, comparing multiple structures simultaneously remains challenging, especially on two-dimensional (2D) displays. Existing 2D visualization tools lack standardized approaches for pipelined inspection of large protein sets, limiting their utility in large-scale pre-filtering.

**Results:** We introduce FlatProt, a tool designed to complement 3D viewers by enabling standardized 2D visualization of individual protein structures or large sets thereof. By including Foldseek-based family rotation alignment or an inertia-based fallback, FlatProt creates consistent and scalable visual representations for user-defined protein structures. It supports domain-aware decomposition, family-level overlays, and lightweight visual abstraction of secondary structures. FlatProt processes proteins efficiently, as showcased on a subset of the human-proteome.

**Conclusion:** FlatProt provides clear, consistent, user-friendly visualizations that support rapid, comparative inspection of protein structures at scale. By bridging the gap between interactive 3D tools and static visual summaries, it enables users to explore conserved features, detect outliers, and prioritize structures for further analysis.

**Availability:** GitHub (https://github.com/t03i/FlatProt); Zenodo (https://doi.org/10.5281/zenodo.15697296).

## Background

Understanding protein structure is important to study biological mechanisms, to guide drug discovery [13, 14], and to annotate molecular function [16]. While three-dimensional (3D) models provide detailed structural insight, experts typically view and analyze these models on two-dimensional (2D) displays; both in interactive tools and static publications. Experts routinely compare protein structures using existing tools allowing them to tap into 3D details. However, as structure datasets grow, the time experts must invest may become limiting. This raises a central question: *how can we generate consistent and scalable 2D representations that support rapid, high-level structural comparisons, and hypothesis generation across many protein structures?*

FlatProt proposes a solution by providing standardized, repeatable 2D visualizations tailored for comparative analysis across protein families or heterogeneous structure sets. It is designed to serve as a visual pre-filter, highlighting shared or divergent features and enabling researchers to prioritize structures for more detailed 3D investigation. The generated static 2D images that can be embedded in non-interactive media or used as compact, icon-like representations of protein structure are reduced to the visual minimum while preserving high-level architectural information.

While 3D viewers such as PyMOL [17], ChimeraX [7], and iCn3D [18] are indispensable for detailed structural analysis, they are not optimized for rapidly repeatable comparison across large structure sets. Generating consistent, publication-ready images often requires extensive manual adjustments of orientation, zoom, rendering styles, and annotations—efforts that must be repeated for each structure and are not easily transferable across proteins with varying domain architecture or length. Scripting can enable automation in some tools, but this typically demands significant user expertise and technical setup. Early workhorses such as MolScript [19] pioneered the generation of high-quality 2D vector graphics from 3D structures, but similarly depend on predefined viewpoints and manual configuration. As a result, producing standardized visual summaries across protein families remains a labor-intensive, iterative task. These tools excel at detailed, expert-guided inspection, but lack built-in support for scalable, standardized, and lightweight visual overviews of heterogeneous structure sets.

To overcome these challenges, several tools have explored 2D representations of protein structures. These include SSDraw [20], ProLego [21], Pro-origami [22], iCn3D [18], and the databases 2DProts [23], and OverProt [24]. While each provides valuable visualization capabilities (Supporting Online Material, Figure S4), they are primarily designed for individual proteins, static domains, or as supplementary features within 3D environments. SSDraw, for instance, flattens proteins into sequential diagrams but lacks support for structural comparison. Pro-origami and ProLego focus on individual fold visualization and are no longer actively maintained. iCn3D offers 2D elements within a powerful 3D viewer but is not designed for bulk export or family-level comparison. 2DProts and OverProt, though extensive, are limited to pre-computed CATH domains [25] and do not support user-defined structures. These constraints make existing 2D tools insufficient for large-scale, customizable, and consistent structure comparisons across diverse datasets.

FlatProt was developed to address this gap. Unlike existing 2D tools, it enables consistent, scalable visualization of user-defined protein sets. By abstracting structural elements into simplified 2D representations and applying standardized orientation schemes, FlatProt facilitates rapid visual comparison while minimizing complexity. Whether used for identifying conserved cores, spotting structural deviations, or preparing non-interactive visuals, generated visualizations act as a structural overview that complements, rather than replaces, detailed 3D inspection. Using FlatProt, researchers are able to quickly screen and compare large structure sets, identify conserved or divergent regions across families, and generate reproducible, standardized visuals for integration into pipelines, figures, or annotation platforms. By bridging the gap between interactive 3D inspection and static high-throughput visualization, FlatProt facilitates scalable structural exploration in both research and communication contexts.

In the following, we introduce the specific capabilities and implementation details of FlatProt and demonstrate the efficacy of FlatProt through several case studies. The FlatProt tool, documentation, example usages, and further information are available through GitHub (https://github.com/t03i/FlatProt).

## Implementation

### Algorithm

FlatProt generates standardized 2D representations of 3D protein structures using a modular pipeline consisting of four core components: *align, project, overlay*, and *split* (Figure 1). These modules enable consistent structure orientation, abstraction of secondary structure, and comparative visualization, while supporting flexible annotations and user-defined styling. The sub-command architecture allows flexible integration in custom workflows, enabling integration into diverse analysis pipelines or selective use of individual features. FlatProt accepts input structures in both PDB and mmCIF formats and integrates optional secondary structure assignments, rotation matrices, and annotations to produce scalable vector graphics (SVG) representations of protein architecture.

**Figure 1:**
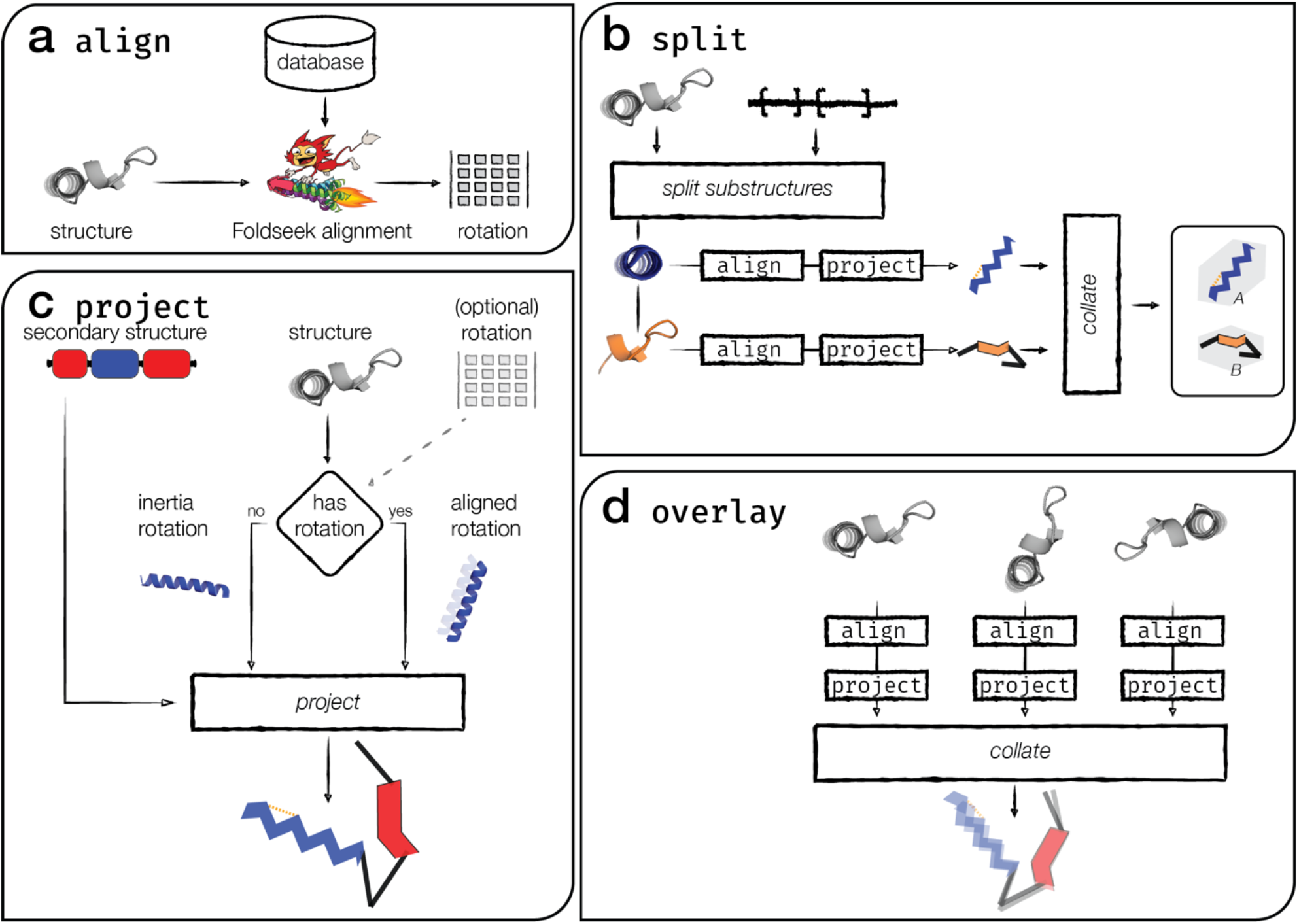
FlatProt - modular visualization pipeline. FlatProt visualizes 3D protein structures through standardized 2D representations using four core modules—align, project, split, and overlay. (a) The align module orients input structures by mapping them to known SCOP superfamilies using Foldseek. If no confident match is found, a fallback inertia-based (PCA) rotation is applied. (b) The split module divides multi-domain structures into annotated segments, aligning and visualizing each independently to reflect domain-specific architecture. (c) The project module transforms each structure into a stylized 2D image, rendering helices as zigzags, strands as triangles, and loops as lines, optionally including annotations such as disulfide bonds or residue labels. (d) The overlay module merges multiple projected structures into a composite image to highlight conserved versus divergent features across families, with opacity encoding structural consensus.

#### Optimal structural alignment through module align

Per default, FlatProt uses the fast structure-alignment method Foldseek [6] with 3Di identity searches to identify the closest family in a pre-generated SCOP Superfamily database [15, 26] (details below: *Family mapping database from SCOP*), which is automatically downloaded on first use. Users may also supply a SCOP Superfamily identifier [15, 26] to return the structural alignment to the respective SCOP domain; alternatively, it is possible to replace the default database with a custom family mapping to match users’ domain definitions or use case. Instructions for generating and using custom databases are provided in the Supporting Online Material (SOM: Custom Database Construction).

If Foldseek returns a confident match, a compatible rotation matrix is calculated and written to the specified output location, where it can subsequently be used for a project command. If no match exceeds the user-defined similarity threshold (default: 0.5), no matrix is produced, and a warning is issued. This threshold follows Foldseek’s general guidance [6] and can be tuned to user needs; 0.5 has yielded robust results in practice, as Foldseek returns the best matching structure.

#### 2D visualization with module project

The *project* module renders the 2D representation for a single protein. It accepts a file with 3D coordinates (PDB or mmCIF), an optional file with secondary structure assignment (required if not embedded in mmCIF), and an optional rotation matrix. When a rotation matrix is available, FlatProt applies given rotations to the atomic coordinates and orthographically projects the 3D structure onto the 2D plane. Without a rotation matrix, FlatProt defaults to an inertia-based orientation (SOM: Algorithm S1). Rotation matrices can be produced through the *align* module or extracted interactively from PyMOL (SOM: Custom Rotation Matrix). An invalid matrix (incorrect formatting or mathematical inconsistencies) triggers a warning, falling back to inertia-based orientation.

FlatProt’s default inertia rotation is consistent with orient commands in tools such as PyMOL, ChimeraX, and X-PLOR [7, 17, 27]. It aligns the principal components of the atomic coordinate distribution such that the largest variance lies along the x-axis, the second along the y-axis, and the smallest along the z-axis, maximizing information in the 2D projection while reducing structural overlap. To illustrate how our inertia-based fallback handles structural extremes, we include two canonical examples: elongated fibrous collagen (UniProtKB P27658 [9-11]) and compact globular hemoglobin (PDB 1A3N [3, 28, 29]) (SOM: Figure S6).

Regular secondary structure elements are simplified: helices appear as zigzag ribbons, strands as filled triangles, the non-regular remainder (often referred to as “loops”) are shown as straight lines between the endpoints of helices and strands. This abstraction deliberately emphasizes clarity and pattern recognition at the expense of fine-grained detail such as loop curvature or strand connectivity.

To support interpretation, *project* optionally indicates N- and C-termini, helix and sheet start– end coordinates, and user-defined annotations. These annotations, e.g., points, lines, or residue-spanning regions, can be supplied in external files and styled using a TOML-based stylesheet format, enabling custom coloring, labels, and graphical nuances (SOM: Custom Annotation & Styling). Elements are layered back-to-front by average z-coordinate value, providing consistent back-to-front depth representation in the 2D image.

#### Multi-structure comparison with module overlay

The *overlay* module combines multiple structures into a single comparative visualization by running *align* and *project* for each input. To reduce clutter in large families, users can optionally apply automatic and customizable Foldseek clustering [6] beforehand. For overlays involving many structures, *overlay* can output raster images (e.g., PNG) instead of SVG to reduce file size and to increase rendering efficiency.

In *overlay* visualizations, alignment consensus is conveyed through opacity: highly conserved structural regions appear darker and more prominent, while divergent regions are more transparent, allowing users to easily detect conserved cores and variable motifs.

#### Domain-specific alignment with module split

The *split* module extends FlatProt’s alignment capabilities to multi-domain proteins. It accepts a structure file and a set of residue-regions (e.g., A:1-30,A:40-60) and splits the input into distinct structural segments. Each segment is independently aligned to its most appropriate family using the *align* module, producing separate rotation matrices per region or falling back to inertia-based if no alignment could be found.

The resulting segments are then projected and visualized into a shared canvas. This approach allows for domain-level resolution of structural similarities and differences; particularly useful when individual domains belong to different superfamilies or vary in conservation. Like other modules, *split* supports annotation and styling, and outputs a single combined SVG.

### Hardware

We performed all tests on commodity hardware to ensure the accessibility and practicality of FlatProt for typical users. Example analyses were performed on a standard Apple MacBook Pro M4, 2025, 36GB RAM.

### Data sets for case studies

*Family mapping database from SCOP*. We created an initial FlatProt database for family mapping from the *SCOP2* database [15, 26] using the superfamily level of protein classification with 2,816 classes. We identified a representative for each superfamily as the structure with the highest average alignment TM-score [30] to all other family members. This yielded 2,783 representatives. Although FlatProt uses SCOP [15] by default, users can preset a custom database, allowing individual adjustments while maintaining family-wide consistency. Alternative sources of family assignments, e.g., CATH [25], can be leveraged to derive a custom database (GitHub repository for more details).

#### Benchmarking performance on the human proteome

We assessed how FlatProt scales with protein length using the human proteome from UniProtKB/SwissProt[9] (data accessed July 2024), with structures predicted from AlphaFoldDB [10, 11]. The dataset initially included 20,420 proteins. We filtered this set by removing 279 proteins with missing coordinates and 814 orphans, yielding 19,327 proteins. Our final evaluation set was constructed by randomly sampling 1,000 proteins (lengths: 68 to 2,671 residues) from this filtered pool, ensuring the sample reflected the overall length distribution and could be aligned to the Family mapping database through the default align command.

#### Kallikreins showcase family visualization

We used the protein family of *Kallikreins* (KLK) to give an example for FlatProt’s visualization. KLK is a subgroup of serine proteases characterized by two attached beta sheets. We took a set of 437 proteins (123-226 residues; mean: 219) identified by experts as family members [4] and predicted 3D structures for each protein using ColabFold [5]. We created a representative subset through clustering with Foldseek [6] (coverage threshold: 0.9, minimal sequence identity: 0.5). This identified 68 clusters, 28 of them singletons (containing only one structure), resulting in 40 family-representative clusters. The predicted structures and representatives are attached in the Supporting Online Material (SOM: KLK.zip).

*3FTx showcases comparison clarity*. We used the protein family of *Three-Finger Toxins (3FTx)* to highlight how FlatProt allows comparisons across many structures. Characterized by a three-finger fold consisting of beta-stranded loops and a central hydrophobic core formed by four conserved disulfide bonds [31], 3FTx proteins vary in the number of cysteine bonds and beta-sheets. Our dataset included 1,427 trimmed structures [32].

## Results and Discussion

### Conserved fold highlights structural variation in 3FTx

To assess whether FlatProt enables interpretable visual comparison across structurally related proteins, we applied it to the Three-finger Toxin (3FTx) family. These proteins share a conserved architecture but include numerous natural variants, making them ideal candidates for visual inspection.

Comparing 3FTx structures in traditional 3D viewers often requires manual reorientation and overlay, which complicates the identification of subtle deviations. In contrast, FlatProt enforces a consistent orientation across all inputs using Foldseek-aligned rotation and a fixed projection scheme. This yields directly comparable, layout-stable images even across structurally diverse members.

The resulting 2D projections (Figure 2) capture the conserved three-finger topology across the family while clearly exposing structural differences. For example, A7X3S2 [8-11] (Figure 2a) displays a shortened and laterally displaced third finger, coinciding with the absence of a disulfide bridge in that region—suggesting a deviation from the canonical fold. Similarly, P01398 [9-12] (Figure 2b) shows an additional helix embedded in one of its fingers, a feature that is prominent in FlatProt’s schematic rendering.

**Figure 2:**
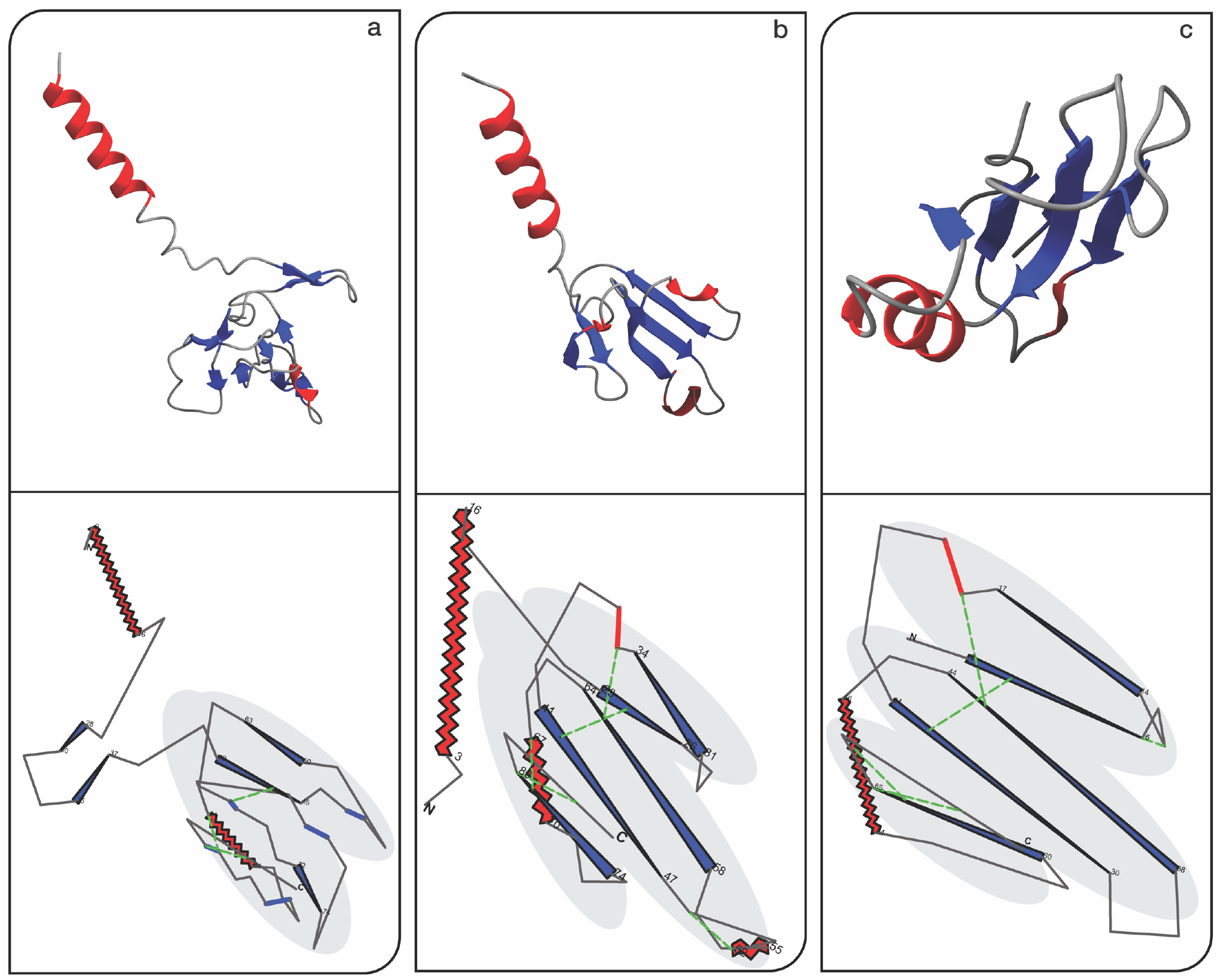
Consistent visualization of 3FTx family: Each panel highlights the structural variations within the Three-finger Toxins (3FTx) family, demonstrating FlatProt’s capability to effectively capture and compare these differences. The core three-finger structure is manually shaded for comparison. To improve interpretability, the upper row shows the corresponding 3D structures of the proteins, rotated using ChimeraX’s [7] orient command for consistency across views. Disulfide bridges are annotated using Chimera’s default bond visualization, but not visible in this perspective. All images were rotated by 90 degrees to align visual axes between 3D and 2D views. (a) A7X3S2 [8-11] (present in Trimorphodon biscutatus) variation within the 3FTx family with a rotated outwards finger in the bottom right; (b) P01398 [9-12] (present in Bungarus multicinctus) a long 3FTx protein (long helix manually rotated for comparability) with a small helix in the second finger; (c) Canonical 3FTx protein with typical three-finger structure (present in Naja Naja [4, 10, 11]).

While FlatProt omits depth cues and fine-grained atomic detail, its orthographic projection emphasizes overall geometry and parallelism, enhancing the visual recognition of structural patterns. Cystine bridges are annotated explicitly [33], and secondary structure elements are rendered as minimalistic shapes to improve readability. To support interpretation, FlatProt also overlays the positions of N- and C-termini and labels the start and end of major secondary structure elements, providing consistent internal orientation markers. These abstractions support fast, scalable comparisons that would be difficult to perform interactively at scale.

Each 3FTx protein was processed in approximately 1.5 seconds, demonstrating FlatProt’s suitability for rapid screening across large protein families. The tool thus enables users to prioritize targets for detailed inspection by highlighting deviations from canonical forms or identifying conserved architectural features.

### Domain splitting clarifies modular architecture in 1KT0

To evaluate FlatProt’s ability to visualize complex, multi-domain proteins, we applied it to the FKBP-like protein 1KT0 [1-3], a 457-residue member of human steroid receptor complexes. These proteins are challenging to interpret in full due to their modular composition, overlapping folds, and variation in domain organization. FlatProt provides both global and domain-resolved views of such proteins (Figure 3). The global projection (Figure 3a), generated via inertia-based rotation, compactly represents the overall shape and structural extent. However, while this view reflects the macro-architecture, it does not clearly delineate internal segmentation or domain boundaries—especially when domains are packed closely or share similar orientation.

**Figure 3:**
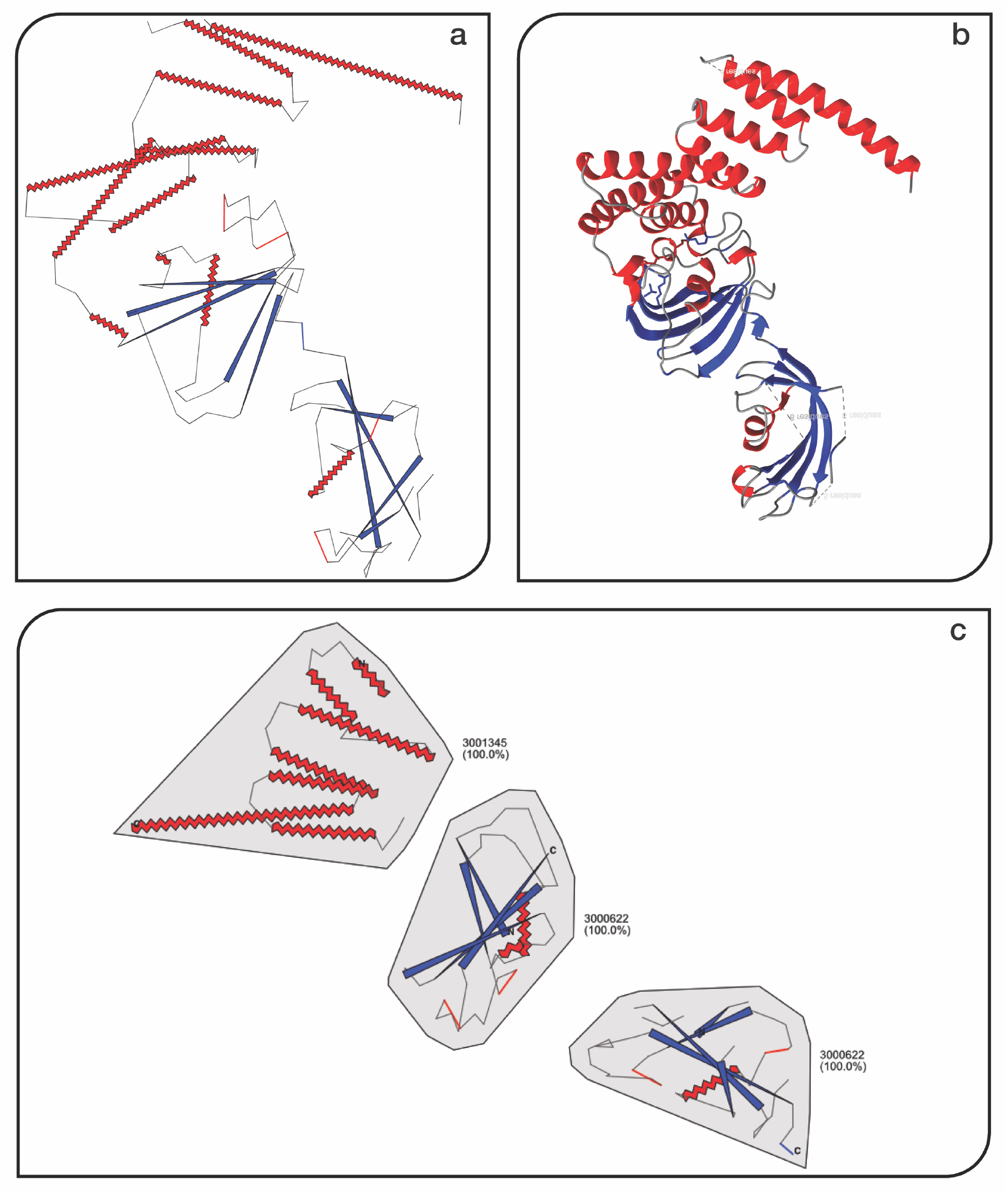
Visualization of 1KT0 [1-3] Protein with Inertia Rotation and Domain Annotations. This figure demonstrates FlatProt’s capability to handle large, multi-domain proteins through domain-aware visualization and consistent orientation. (a) FlatProt visualization of 1KT0 [1-3] using the inertia rotation algorithm, highlighting the global architecture without domain eparation. (b) Corresponding 3D structure of 1KT0 exported from ChimeraX [7] after applying the initial orientation. The mage has been mirrored horizontally to align with the right-handed coordinate system of the 2D projection in panel (a), ensuring visual consistency. (c) Domain-level visualization using Chainsaw-predicted segmentation, with each domain ndependently aligned to its most appropriate family and visualized using FlatProt.

To increase interpretability, we applied FlatProt’s domain splitting feature using predicted segmentation from Chainsaw [34]. Each domain was processed and aligned independently using Foldseek [6] (Figure 3c). The resulting visualization reveals a clear separation between domains and highlights structural duplication: both the first and second segments were assigned to the same SCOP superfamily (3000622) [15], with visually consistent layout and scale. These repeated domains become immediately apparent in the split view, while they remain difficult to distinguish in the full-length projection. Although secondary structures such as beta sheets are abstracted and not aligned across domains, the standardized rendering of helices, strands, and loop elements supports structural comparison at a high level. FlatProt overlays orientation markers, including N- and C-terminal labels, aiding visual navigation across the split layout.

This example illustrates how FlatProt’s modular processing enhances interpretability for large proteins by enabling both overview and detailed inspection. It also demonstrates the tool’s utility in highlighting domain-level redundancy, even in cases where 3D superposition or sequence alone may obscure structural repetition.

### Visualizing conserved family features through structure overlays

FlatProt’s overlay mode enables direct visual comparison of structural conservation across large protein families. To assess its ability to reveal consensus features, we applied it to the Kallikrein (KLK) family, a group of serine proteases with well-characterized domain structure but substantial variation across homologs [4, 5].

In conventional 3D viewers, overlaying many structures often results in visual clutter or requires manual filtering and adjustment to highlight conserved regions. FlatProt instead uses automated clustering and representative sampling to generate a concise overlay. By projecting each selected structure into a shared reference frame, it visually reinforces recurring architectural elements while de-emphasizing rare or noisy deviations. The KLK overlay (Figure 4) reveals a stable core of aligned structural elements, with increased opacity indicating recurring motifs across the representatives. Notably, two helical elements appear in a substantial subset of the family, giving rise to prominent, high-opacity bands. While FlatProt does not resolve beta-sheet topology directly, repeated strand positions become visible through their consistent orientation and placement in the overlay.

**Figure 4:**
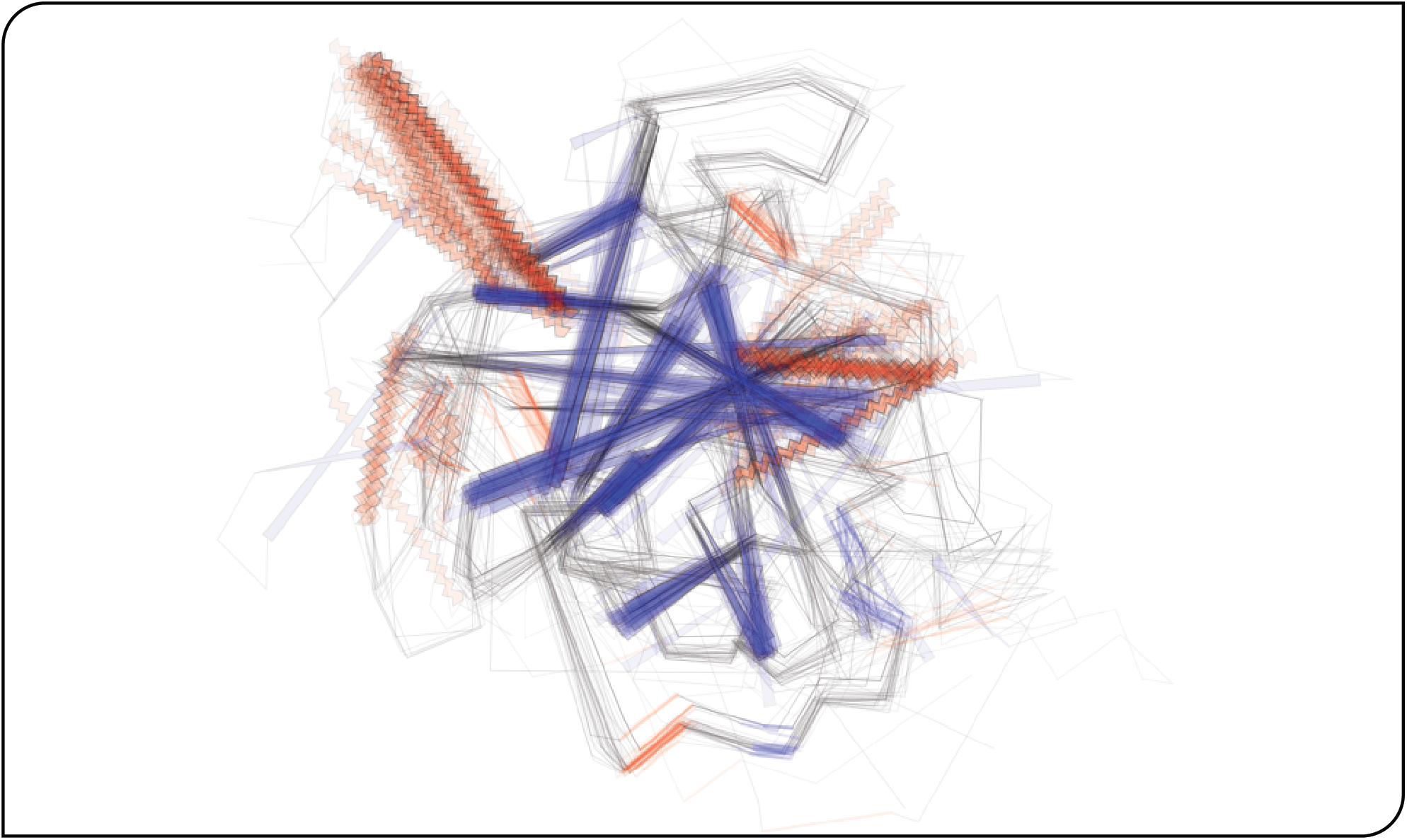
Structural overlay of KLK family representatives. Shown are 40 representative structures of the Kallikrein (KLK) family [4, 5], selected through Foldseek [6] clustering and visualized using FlatProt. All structures are aligned to the SCOP superfamily 3000114 [15] to ensure consistent orientation. The resulting 2D projections are overlaid to highlight conserved versus variable regions: highly conserved substructures appear with high opacity and sharp contours, while lower-opacity traces indicate divergent or infrequent features. A full 3D structural overlay is provided in the Supporting Online Material (Figure S2).

Compared to full-resolution 3D viewers, FlatProt sacrifices fine detail—such as side-chain packing or strand connectivity—to emphasize large-scale architectural trends. This trade-off allows conserved structural features to stand out more clearly and reduces distraction from unique, low-frequency deviations. The result is a compact, layout-consistent visualization that supports intuitive interpretation. Researchers can rapidly assess which features are broadly conserved, detect potential subfamily-specific insertions, and identify structurally deviant representatives.

### Scalable performance through integrated alignment and fast projection

To assess FlatProt’s suitability for large-scale applications, we benchmarked 1,000 human proteins that yielded successful Foldseek family alignments. Each structure was processed in two workflows: first using *align+project* for family-consistent orientation and again using *project* with an inertia-based fallback (Figure 5).

**Figure 5:**
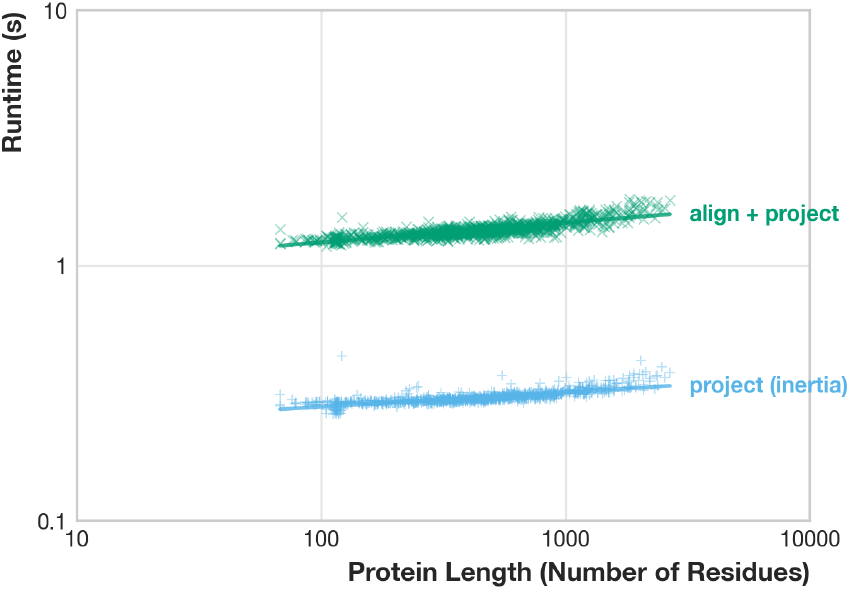
Runtime scaling of common FlatProt workflows on human proteins. This log-log scatter plot shows runtimes for project (inertia-based fallback, blue) and align+project (Foldseek-based family alignment, green) across 1,000 proteins from the human proteome. Inertia runtimes scale with protein length (slope = 0.06, R^2^ = 0.61) and remain between 0.26–0.44 s. Foldseek-aligned projections are slower but less length-dependent (slope = 0.08, R^2^ = 0.68), completing in 1.19–1.83 s. Output file sizes scale sub-quadratically and remain below 1.4 MB (SOM: Figure S5).

In the alignment-based workflow, FlatProt uses Foldseek to identify the closest SCOP superfamily and applies the corresponding rotation matrix before projecting the structure. This ensures that related proteins are rendered with consistent orientation, which is critical for family overlays and comparative inspection.

In the fallback workflow, FlatProt orients structures solely based on their geometry. It computes the inertia tensor of the atomic coordinates and aligns the principal axes to the 2D plane. This approach is fast and robust and requires no external reference or alignment.

The results indicate that inertia-based projections complete in well under 0.5 s per structure and scale predictably with protein length. Family-aligned projections take slightly longer (1.2–1.8 s), but their runtimes are largely independent of sequence length due to Foldseek’s optimization. Importantly, both workflows produce SVG files that remain lightweight even for long proteins, enabling fast rendering and efficient storage (SOM: Figure S5).

These benchmarks confirm that FlatProt balances speed, scalability, and alignment fidelity. Its fallback mode supports rapid standalone inspection, while the alignment-integrated workflow enables consistent visualization across large, diverse structure sets.

## Conclusion

The rapid growth of predicted protein structures—driven by resources like AlphaFoldDB—has created new demands for scalable, interpretable structural comparison. While interactive 3D tools remain essential for detailed analysis, they are ill-suited for high-throughput workflows, static communication formats, or intuitive pattern recognition across large datasets.

FlatProt addresses this gap by introducing a new modality of structural visualization: one that is scalable, standardized, and minimal. Its non-interactive, 2D output enables consistent layout, rapid inspection, and integration into publications, annotation pipelines, or automated screens. By abstracting complex 3D structures into simplified visual forms, FlatProt allows researchers to visually filter and prioritize proteins across families—before engaging with full 3D inspection.

Case studies across diverse proteins demonstrate FlatProt’s ability to reveal conserved folds, detect variation, and clarify domain structure. The combination of fast inertia-based projection and Foldseek-guided alignment ensures both speed and contextual comparability. Output files remain compact and interpretable, even at proteome scale.

FlatProt deliberately trades fine-grained detail for visual clarity and standardization. While atom-level features and depth cues are omitted, the result is a visual pre-filter that enhances structural intuition and supports large-scale exploration. It empowers users not only to inspect proteins, but to generate insight and communicate structure-function relationships with minimal overhead.

Looking ahead, we aim to extend FlatProt with a more expressive 2D design language tailored to protein engineering and visualization, to improve the alignment of higher-order secondary structure patterns such as beta-sheets, and to provide a web-based interface that broadens accessibility. As structural biology continues to scale in complexity, tools like FlatProt will be essential—not just for inspection, but for comparative reasoning, intuitive discovery, and effective communication.

## Supporting information

KLK.zip

SOM

## Abbreviations

3D: three-dimensional
2D: two-dimensional
DSSP: Dictionary of Secondary Structure of Proteins (automatic assignment of secondary structure from 3D co-ordinates)
KLK: Kallikreins
3FTx: Three-Finger Toxins
ID: Identifier
LDDT: Local Distance Difference Test
mmCIF: Macromolecular Crystallographic Information File (file format)
PDB: the Protein Data Bank (database of 3D coordinates of high-resolution experimental structures
PNG: Portable Network Graphics (file format)
SCOP: Structural Classification of Proteins
SVG: Scalable Vector Graphics (file format)
UniProtKB: Universal Protein Resource Knowledgebase (database).

## Availability and requirements

**Project name:** FlatProt

**Project home page:** https://github.com/t03i/FlatProt

**Operating system(s):** Platform independent

**Programming language:** Python 3.12.5

**Other requirements:** python version “>=3.10, <3.13”, drawsvg>=2.4.0, gemmi>=0.7.0, polars>=1.17.1, pydantic-extra-types>=2.10.1, pydantic>=2.10.3, pydantic-settings>=2.6.1, tqdm>=4.67.1, cyclopts>=3.9.0, toml>=0.10.2, h5py>=3.13.0, numpydantic>=1.6.8, platformdirs>=4.3.6, numpy>=2.1.3, Foldseek, DSSP (mkdssp version >=4.2.2), Cairo

**License:** Apache 2.0

**Any restrictions on use by non-academics:** None

## Declarations

### Availability of data and materials

This published article and its supplementary information files include all data generated or analyzed during this study. The generated database is available on Zenodo https://doi.org/10.5281/zenodo.15697296

### Authors’ contributions

**Tobias Olenyi:** Conceptualization, Software, Methodology, Visualization, Writing - Original Draft, Writing - Review & Editing, Project Administration

**Constantin Carl:** Software, Methodology, Formal Analysis, Writing - Original Draft,

**Tobias Senoner:** Data Curation, Methodology, Writing - Review & Editing, Project Administration

**Ivan Koludarov:** Conceptualization, Supervision, Data Curation, Writing - Review & Editing. Funding Acquisition

**Burkhard Rost:** Supervision, Writing - Review & Editing, Funding Acquisition

### Declaration of generative AI and AI-assisted technologies in the writing process

While preparing this work, the authors used Claude.ai and ChatGPT to check the language and grammar of this manuscript. After using this service, the authors reviewed and edited the content as needed and take full responsibility for the content of the published article.

## Acknowledgments

We thank Nikita Kugut (TUM) for his support with many aspects of this work. We thank the Technical University of Munich (TUM) for providing facilities and resources. We are also grateful to Alex Stivala, the creator of Pro-origami, for maintaining a website with related protein 2D visualization tools, as well as to all contributors to open-source programming libraries. Finally, we thank those who deposit experimental data in public databases, maintain these databases, and develop methods to enrich experimental data.

## References

1. Sinars CR, Cheung-Flynn J, Rimerman RA, Scammell JG, Smith DF, Clardy J: Structure of the large FK506-binding protein FKBP51, an Hsp90-binding protein and a component of steroid receptor complexes. Proc Natl Acad Sci USA 2003, 100(3):868–873.

2. Sinars CR, Cheung-Flynn J, Rimerman RA, Scammell JG, Smith DF, Clardy JC: 2003.

3. Berman HM, Westbrook J, Feng Z, Gilliland G, Bhat TN, Weissig H, Shindyalov IN, Bourne PE: The Protein Data Bank. Nucleic Acids Res 2000, 28(1):235–242.

4. Barua A, Koludarov I, Mikheyev AS: Co-option of the same ancestral gene family gave rise to mammalian and reptilian toxins. BMC Biol 2021, 19(1):268.

5. Mirdita M, Schutze K, Moriwaki Y, Heo L, Ovchinnikov S, Steinegger M: ColabFold: making protein folding accessible to all. Nat Methods 2022, 19(6):679–682.

6. Van Kempen M, Kim SS, Tumescheit C, Mirdita M, Lee J, Gilchrist CLM, Söding J, Steinegger M: Fast and accurate protein structure search with Foldseek. Nat Biotechnol 2024, 42(2):243–246.

7. Meng EC, Goddard TD, Pettersen EF, Couch GS, Pearson ZJ, Morris JH, Ferrin TE: UCSF ChimeraX: Tools for structure building and analysis. Protein Science 2023, 32(11):e4792.

8. Fry BG, Scheib H, van der Weerd L, Young B, McNaughtan J, Ramjan SF, Vidal N, Poelmann RE, Norman JA: Evolution of an arsenal: structural and functional diversification of the venom system in the advanced snakes (Caenophidia). Mol Cell Proteomics 2008, 7(2):215–246.

9. The UniProt C, Bateman A, Martin M-J, Orchard S, Magrane M, Ahmad S, Alpi E, Bowler-Barnett EH, Britto R, Bye-A-Jee H et al: UniProt: the Universal Protein Knowledgebase in 2023. Nucleic Acids Research 2023, 51(D1):D523–D531.

10. Jumper J, Evans R, Pritzel A, Green T, Figurnov M, Ronneberger O, Tunyasuvunakool K, Bates R, Zidek A, Potapenko A et al: Highly accurate protein structure prediction with AlphaFold. Nature 2021, 596(7873):583–589.

11. Varadi M, Bertoni D, Magana P, Paramval U, Pidruchna I, Radhakrishnan M, Tsenkov M, Nair S, Mirdita M, Yeo J et al: AlphaFold Protein Structure Database in 2024: providing structure coverage for over 214 million protein sequences. Nucleic Acids Res 2024, 52(D1):D368–D375.

12. Grant GA, Chiappinelli VA: kappa-Bungarotoxin: complete amino acid sequence of a neuronal nicotinic receptor probe. Biochemistry 1985, 24(6):1532–1537.

13. Lv Q, Chen G, Yang Z, Zhong W, Chen CY: Meta-MolNet: A Cross-Domain Benchmark for Few Examples Drug Discovery. IEEE Trans Neural Netw Learn Syst 2025, 36(3):4849–4863.

14. Lv Q, Chen G, Yang Z, Zhong W, Chen CY: Meta Learning With Graph Attention Networks for Low-Data Drug Discovery. IEEE Trans Neural Netw Learn Syst 2024, 35(8):11218–11230.

15. Chandonia J-M, Guan L, Lin S, Yu C, Fox Naomi K, Brenner Steven E: SCOPe: improvements to the structural classification of proteins – extended database to facilitate variant interpretation and machine learning. Nucleic Acids Research 2022, 50(D1):D553–D559.

16. O’Donoghue SI, Sabir KS, Kalemanov M, Stolte C, Wellmann B, Ho V, Roos M, Perdigão N, Buske FA, Heinrich J et al: Aquaria: simplifying discovery and insight from protein structures. Nat Methods 2015, 12(2):98–99.

17. Schrödinger LLC: The PyMOL Molecular Graphics System, Version 1.8. In.; 2015.

18. Wang J, Youkharibache P, Marchler-Bauer A, Lanczycki C, Zhang D, Lu S, Madej T, Marchler GH, Cheng T, Chong LC et al: iCn3D: From Web-Based 3D Viewer to Structural Analysis Tool in Batch Mode. Front Mol Biosci 2022, 9:831740.

19. Kraulis PJ: MOLSCRIPT: a program to produce both detailed and schematic plots of protein structures. Journal of Applied Crystallography 1991, 24(5):946–950.

20. Chen EA, Porter LL: SSDraw: software for generating comparative protein secondary structure diagrams. In.; 2023.

21. Khan T, Panday SK, Ghosh I: ProLego: tool for extracting and visualizing topological modules in protein structures. BMC Bioinformatics 2018, 19(1):167.

22. Stivala A, Wybrow M, Wirth A, Whisstock JC, Stuckey PJ: Automatic generation of protein structure cartoons with Pro-origami. Bioinformatics 2011, 27(23):3315–3316.

23. Hutařová Vařeková I, Hutař J, Midlik A, Horský V, Hladká E, Svobodová R, Berka K: 2DProts: database of family-wide protein secondary structure diagrams. Bioinformatics 2021, 37(23):4599–4601.

24. Midlik A, Hutarova Varekova I, Hutar J, Chareshneu A, Berka K, Svobodova R: OverProt: secondary structure consensus for protein families. Bioinformatics 2022, 38(14):3648–3650.

25. Waman VP, Bordin N, Lau A, Kandathil S, Wells J, Miller D, Velankar S, Jones DT, Sillitoe I, Orengo C: CATH v4.4: major expansion of CATH by experimental and predicted structural data. Nucleic Acids Res 2025, 53(D1):D348–D355.

26. Fox NK, Brenner SE, Chandonia J-M: SCOPe: Structural Classification of Proteins—extended, integrating SCOP and ASTRAL data and classification of new structures. Nucleic Acids Research 2014, 42(D1):D304–D309.

27. Badger J, Kumar RA, Yip P, Szalma Sn: New features and enhancements in the X-PLOR computer program. Proteins: Structure, Function, and Genetics 1999, 35(1):25–33.

28. Tame J, Vallone B: DEOXY HUMAN HEMOGLOBIN - 1A3N. 1998.

29. Tame JR, Vallone B: The structures of deoxy human haemoglobin and the mutant Hb Tyralpha42His at 120 K. Acta Crystallogr D Biol Crystallogr 2000, 56(Pt 7):805–811.

30. Zhang Y, Skolnick J: Scoring function for automated assessment of protein structure template quality. Proteins 2004, 57(4):702–710.

31. Kini RM, Doley R: Structure, function and evolution of three-finger toxins: Mini proteins with multiple targets. Toxicon 2010, 56(6):855–867.

32. Koludarov I, Senoner T, Jackson TNW, Dashevsky D, Heinzinger M, Aird SD, Rost B: Domain loss enabled evolution of novel functions in the snake three-finger toxin gene superfamily. Nat Commun 2023, 14(1):4861.

33. Sun M-a, Wang Y, Zhang Q, Xia Y, Ge W, Guo D: Prediction of reversible disulfide based on features from local structural signatures. BMC Genomics 2017, 18(1):279.

34. Wells J, Hawkins-Hooker A, Bordin N, Sillitoe I, Paige B, Orengo C: Chainsaw: protein domain segmentation with fully convolutional neural networks. In.; 2023.

