## Supplementary material for "FlatProt: 2D visualization eases protein structure comparison": SOM

### FlatProt: 2D visualization eases protein structure comparison - Supporting Material

#### Contents

This supplementary material provides technical details and extended benchmarking for FlatProt, a tool designed for standardized 2D visualization of protein structures. It outlines the fallback inertia-based rotation algorithm in pseudocode, presents ChimeraX-based 3D overlays for structural validation, and compares FlatProt's performance with alternative tools like PyMOL, ChimeraX, SSDraw, and Pro-origami. The document includes visual comparisons of 2D projections from different software, analysis of SVG file size scaling with protein length, and demonstrations of FlatProt's ability to distinguish structural classes such as globular and fibrous proteins. Additionally, it documents user-configurable features, including custom annotation, styling via TOML, manual matrix extraction for orientation control, and the construction of custom alignment databases.

#### Inertia-Based Rotation Algorithm in Pseudo-Code

<u>Inertia-Based Rotation</u>$(coordinates)$:

// Compute center of geometry and center
let $C = \textbf{mean}(\text{coordinates})$
let $X = \text{coordinates} - C$

// Compute Inertia Tensor for all coordinates
let $I = \textbf{zero-matrix}(3, 3)$
for $r$ in $X$:
    let $r_{\text{squared}} = \textbf{dot}(r, r)$
    let $I = I + \left(r_{\text{squared}} \cdot \textbf{identity}(3) - \textbf{outer}(r, r)\right)$

// Eigendecomposition of Inertia Matrix
let $(\text{eigenvalues}, \text{eigenvectors}) = \textbf{eigendecompose}(I)$
let $R = \text{eigenvectors}$

// Ensure right-handed system
if $\textbf{determinant}(R) < 0$:
    $R_{\{[:,2]\}} = -R_{\{[:,2]\}}$

// Rotate and Translate
let $T = -R \cdot C$
let $X_{\text{aligned}} = \textbf{transpose}(R \cdot \textbf{transpose}(\text{coordinates}) + T)$
return $X_{\text{aligned}}$

**Algorithm S1**: Inertia-based rotation algorithm used by FlatProt as fallback if no rotation matrix is available.

#### Structure Overlay in ChimeraX

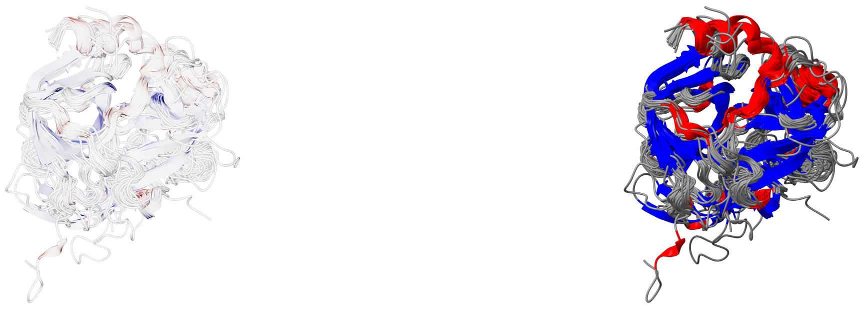

(a) Overlay with each structure at 20% opacity. (b) Overlay with each structure at full opacity.

**Figure S2**: **Overlay of 40 KLK family structures using ChimeraX MatchMaker alignment**. This figure shows two comparative overlays, generated using ChimeraX's MatchMaker algorithm [1], applied to 40 representative KLK structures. All models are displayed as cartoon representations and were aligned to `data_Alligator_sinensis_GeneID=102372276.cif`, the first structure in the list. **Figure** 2a shows each structure rendered at 20% opacity. While this highlighting reveals some overlap, the non-additive transparency causes conserved regions to appear only faintly. In contrast, **Figure** 2b displays all structures at full opacity, revealing unmatched structural detail and precise alignment. Determining conserved regions requires a trained eye to identify areas of tight structural superposition, which indicate evolutionary conservation. These overlays underscore the trade-off between interpretability and completeness in 3D superpositions, and complement FlatProt's minimal, standardized projections by preserving atom-level resolution and spatial context.

#### Performance Comparison Alternative Methods

| Tool | Single | Family | Family Large |
|---|---|---|---|
| Pro-origami [2] | 250.2±2.4s | – | – |
| SSDraw [3] | 2.298±0.017s | – | – |
| **FlatProt** | 0.265±0.000s<br>(1.361±0.001s) | 1.711±0.005s | 4.156±0.023s |
| ChimeraX [1] | 2.408±0.009s | 2.702±0.008s | 15.041±0.049s |
| PyMol [4] | 0.215±0.001s | 3.376±0.019s | 120.25±0.27s |

**Table S3**: **Runtime comparison of structure visualization tools.** This table compares the average runtime (± standard error) of different tools when generating 2D visualizations of KLK protein structures across three benchmarking tasks. Each task was repeated 10 times per structure or operation to ensure statistical robustness.

- **Single:** Each tool was used to generate image representations for 40 KLK representative structures individually. For FlatProt, the first time shown corresponds to the PCA-based "inertia rotation," while the bracketed time reflects the combined duration of family alignment and rotation using a transformation matrix.
- **Family:** Tools were tasked with generating a family overlay of the 40 KLK representative structures. PyMOL and ChimeraX aligned all structures to the first one (cf. **Figure** 2b), while FlatProt aligned each structure to a family-wide consensus. SSDraw and Pro-origami do not support structural overlays and are marked accordingly.
- **Family Large:** Tools attempted to visualize overlays of all 436 KLK family members. FlatProt used internal clustering to reduce the number of structures and simplify the overlay, whereas PyMOL and ChimeraX processed the full, unfiltered set, resulting in higher runtimes.

FlatProt was faster than any other tool except PyMOL in the single-image generation task, even when including alignment, and consistently outpaces all tools in the overlay tasks. All tests were conducted on a MacBook M4 Pro, except for Pro-origami, which had to be run on an Intel-based 2021 MacBook Pro due to compatibility constraints. This significantly increased its runtime due to the need for multiple emulation and compatibility layers.

#### Visual Comparison of 2D Representations by Different Tools

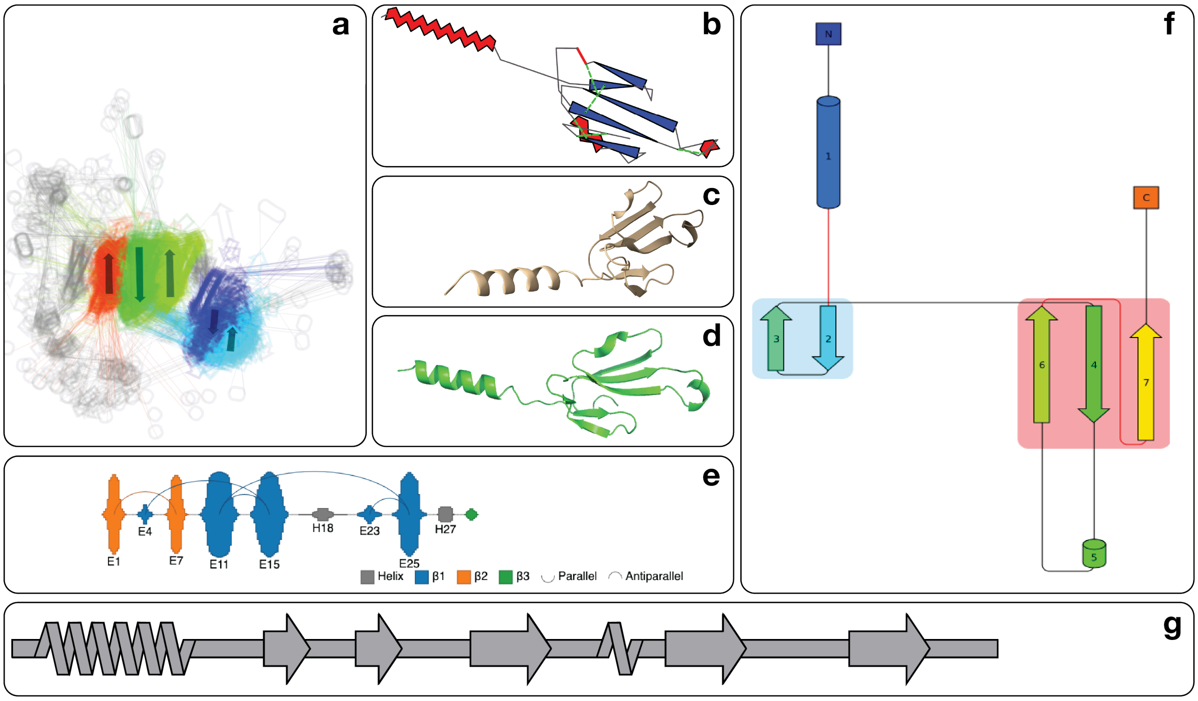

**Figure S4**: **Comparison of 2D structure representations for the 3FTx protein A7X3S2 [5–7] from** Trimorphodon biscutatus**.** These panels illustrate how different tools depict the same protein structure, highlighting variation in representation styles and consistency.

**(a)** Representation of the CATH family 2.10.60.10 [8] in Prots2D, annotated by structural family [9]. This image was manually retrieved, as automated access was not possible. **(b)** FlatProt projection of A7X3S2, including disulfide bridge annotations in green. **(c)** ChimeraX depiction with the structure oriented for maximal inertia in the x,y plane [1]. **(d)** PyMOL rendering using PCA-based orientation, maximizing variance in the x- and y-dimensions [4]. **(e)** CATH family 2.10.60.10 [8] as shown in OverProt [10]. This image was also retrieved manually due to lack of batch export support. **(f)** Pro-origami schematic of the structure [2]. **(g)** SSDraw representation of A7X3S2 based on its secondary structure elements [3]. *Note: A secondary structure cartoon from iCn3D [11] could not be retrieved, as the feature was inaccessible during the evaluation period. ProLego [12] was also unavailable at the time of testing.*

The heterogeneity of individual panels highlights the specific use-cases each tool is optimized for. Compared to panels **(a)**, **(e)**, **(f)**, and **(g)**, FlatProt **(b)** emphasizes structural fidelity more directly. In contrast with the 3D-derived projections in **(c)** and **(d)**, FlatProt's clean lines and absence of shading enhance suitability for standardized 2D presentations.

#### File Size in Comparison to Protein Sequence Length

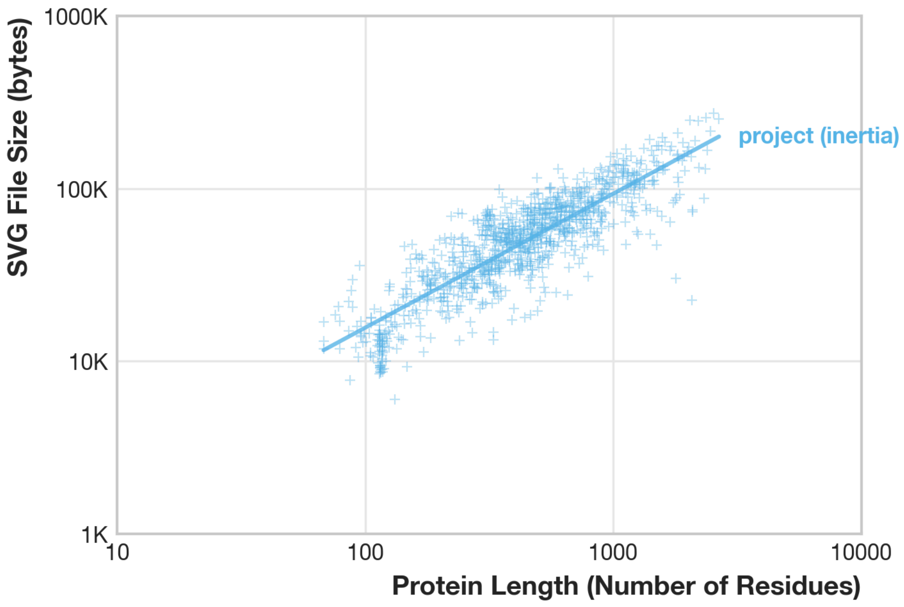

**Figure S5**: **Comparison of Produced FlatProt Projection Output vs. Protein Length** This log-log scatter plot shows the relationship between protein length (in residues) and SVG file size (in bytes) for 1000 protein structures rendered using the FlatProt projection with default inertia-based rotation. The file size grows according to a sub-quadratic power-law with protein length ($\log_{10}(\text{size}) = 0.78 \times \log_{10}(\text{length}) + 2.64, R^2 = 0.74$), indicating consistent but slower-than-quadratic growth. Even for very large proteins—e.g., 30,000 residues, the estimated file size remains below 1.4 MB, which is still manageable for rendering and interaction on most modern machines.

#### Inertia-Based Rotation Captures Fibrous vs. Globular Structures

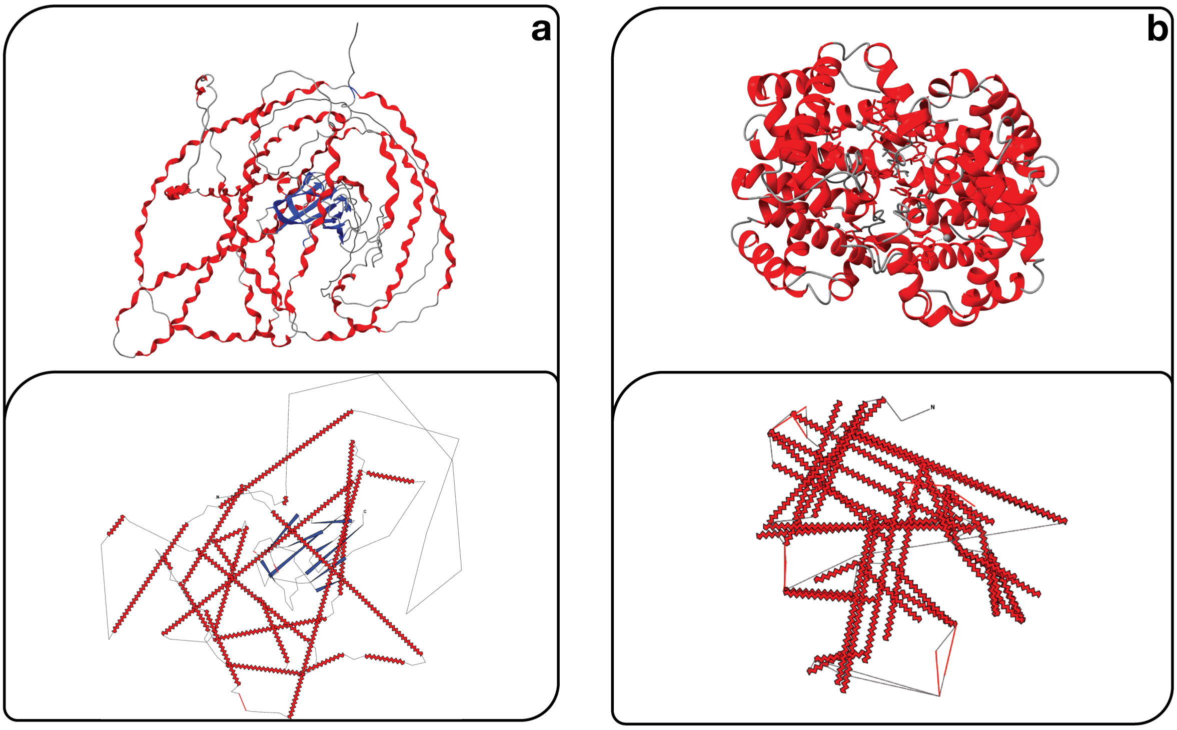

**Figure S6**: **Comparison of Inertia-Based Projection for Fibrous and Globular Proteins** FlatProt's inertia-based rotation captures structural extremes, offering consistent, interpretable 2D layouts across diverse protein classes. **(a)** *Collagen alpha-1(VIII) chain* (UniProt: P27658 [5–7]), a fibrous AlphaFold2-predicted protein, is shown in ChimeraX (top, orient command) and in FlatProt (bottom). While ChimeraX respects atomic realism, FlatProt's projection prioritizes high-level geometry: the elongated, helical architecture is preserved, emphasizing the protein's fibrous character. **(b)** *Hemoglobin complex* (PDB: 1A3N [13–15]), a tetrameric globular protein, is shown using ChimeraX (top) and FlatProt (bottom). Despite its quaternary complexity, FlatProt collapses symmetry-related chains into a simplified 2D representation by projecting the structure into a shared principal axis frame. This deduplicates symmetric subunits and yields a compact, canonical layout of the dimer-of-dimers architecture. Together, these cases illustrate how FlatProt's inertia-based projection preserves dominant geometric features while reducing structural redundancy—enabling clear, comparative 2D views across protein types.

#### Custom Annotation & Styling

In FlatProt, custom annotations are defined using two external TOML files for maximum flexibility. The primary *annotation file* is used to specify features like points, lines, or residue-spanning regions. For fine-grained control, styling for individual annotations can be defined directly inline within this file. To ensure a consistent look and feel of secondary structure elements, a separate *style sheet* is used to configure the global appearance, defining default visual attributes—such as color, labels, and transparency—for secondary structure elements.

Detailed specifications for the annotation file format and the style file format are available online. A concrete example of this workflow is demonstrated in our dedicated Colab notebook and further illustrated in our general examples.

#### Custom Database Construction

FlatProt supports the creation of user-defined structural alignment databases, enabling standardized visualization and comparison of protein families beyond the scope of the default SCOP-derived dataset. This functionality is particularly useful for analyzing non-canonical or proprietary protein sets, assessing specific structural subclasses, or ensuring consistent orientation across a custom-defined family. The custom database creation pipeline generates a fully functional FlatProt-compatible database directory comprising:

(a) an HDF5 alignment database (`alignments.h5`) containing rotation matrices and metadata,
(b) a Foldseek-compatible search database (`foldseek/`), and
(c) a machine-readable metadata file (`database_info.json`) for validation and reproducibility.

To ensure optimal alignment quality and visual comparability, users are encouraged to manually orient their structures before database construction using tools such as PyMOL [4] or ChimeraX [1]. This manual rotation can then be preserved during database generation, bypassing automatic reorientation steps. The full workflow, including usage examples, input requirements, transformation guidelines, and Foldseek integration, is documented at: https://t03i.github.io/FlatProt/tools/custom_database/.

#### Manual Interactive Matrix Extraction

FlatProt supports custom orientation matrices to enforce user-defined projections. The provided `get_matrix.py` script enables interactive orientation of protein structures in PyMOL [4] and extracts the resulting transformation matrix for use with FlatProt `project`.

After rotating the structure to the desired view, the script captures PyMOL's internal camera matrix, converts it to object space via matrix inversion, applies a Y-axis correction to resolve coordinate system differences, and exports the result as `rotation_matrix.npy`, a 4×3 matrix, compatible with FlatProt.

This workflow allows precise manual control of structural orientation to account for user-specific needs. Full usage instructions are available at: https://t03i.github.io/FlatProt/tools/matrix_extraction/

#### Description *SOM-KLK.zip*

This archive contains structure data used for the family overlay visualization in Figure 4 of the main manuscript. All individual structures used in the analysis are located in the `structures/` subfolder. The `representative_structures/` subfolder contains a curated subset of 40 representative structures selected using `foldseek easy-cluster --min-seq-id 0.5 -c 0.9`. Secondary structure annotations were computed using `mkdssp v. 4.4.10` [16]. These representatives were aligned against the FlatProt database and projected into a shared coordinate frame. A combined structural 3D overlay of the representative structure, created with ChimeraX [1], is displayed in **Figure S2**.

#### SOM Bibliography

1. Meng EC, Goddard TD, Pettersen EF, et al (2023) UCSF ChimeraX: Tools for structure building and analysis. Protein Science 32:. https://doi.org/10.1002/pro.4792
2. Stivala A, Wybrow M, Wirth A, et al (2011) Automatic generation of protein structure cartoons with Pro-origami. Bioinformatics 27:3315–3316. https://doi.org/10.1093/bioinformatics/btr575
3. Chen EA, Porter LL (2023) SSDraw: software for generating comparative protein secondary structure diagrams. https://doi.org/10.1101/2023.08.25.554905
4. Schrödinger, LLC (2015) The PyMOL Molecular Graphics System, Version 1.8
5. Bateman A, Martin M-J, Orchard S, et al (2022) UniProt: the Universal Protein Knowledgebase in 2023. Nucleic Acids Research 51:D523–D531. https://doi.org/10.1093/nar/gkac1052
6. Jumper J, Evans R, Pritzel A, et al (2021) Highly accurate protein structure prediction with AlphaFold. Nature 596:583–589. https://doi.org/10.1038/s41586-021-03819-2
7. Varadi M, Bertoni D, Magana P, et al (2023) AlphaFold Protein Structure Database in 2024: providing structure coverage for over 214 million protein sequences. Nucleic Acids Research 52:D368–D375. https://doi.org/10.1093/nar/gkad1011
8. Sillitoe I, Bordin N, Dawson N, et al (2020) CATH: increased structural coverage of functional space. Nucleic Acids Research 49:D266–D273. https://doi.org/10.1093/nar/gkaa1079
9. Hutařová Vařeková I, Hutař J, Midlik A, et al (2021) 2DProts: database of family-wide protein secondary structure diagrams. Bioinformatics 37:4599–4601. https://doi.org/10.1093/bioinformatics/btab505
10. Midlik A, Hutařová Vařeková I, Hutař J, et al (2022) OverProt: secondary structure consensus for protein families. Bioinformatics 38:3648–3650. https://doi.org/10.1093/bioinformatics/btac384
11. Wang J, Youkharibache P, Marchler-Bauer A, et al (2022) iCn3D: From Web-Based 3D Viewer to Structural Analysis Tool in Batch Mode. Frontiers in Molecular Biosciences 9:. https://doi.org/10.3389/fmolb.2022.831740
12. Khan T, Panday SK, Ghosh I (2018) ProLego: tool for extracting and visualizing topological modules in protein structures. BMC Bioinformatics 19:. https://doi.org/10.1186/s12859-018-2171-9
13. Tame J, Vallone B (1998) DEOXY HUMAN HEMOGLOBIN. http://doi.org/10.2210/pdb1a3n/pdb
14. Tame JRH, Vallone B (2000) The structures of deoxy human haemoglobin and the mutant Hb Tyrα42His at 120 K. Acta Crystallographica Section D Biological Crystallography 56:805–811. https://doi.org/10.1107/s0907444900006387
15. Berman HM (2000) The Protein Data Bank. Nucleic Acids Research 28:235–242. https://doi.org/10.1093/nar/28.1.235
16. Joosten RP, Beek TAH te, Krieger E, et al (2010) A series of PDB related databases for everyday needs. Nucleic Acids Research 39:D411–D419. https://doi.org/10.1093/nar/gkq1105
